# The individual determinants of morning dream recall

**DOI:** 10.1101/2024.05.23.595531

**Authors:** Valentina Elce, Damiana Bergamo, Giorgia Bontempi, Bianca Pedreschi, Michele Bellesi, Giacomo Handjaras, Giulio Bernardi

**Author notes:** **Correspondence:** Giulio Bernardi, IMT School for Advanced Studies Lucca, Piazza San Francesco, 19, 55100 Lucca – Italy.

## Abstract

Evidence suggests that (almost) everyone dreams during their sleep and may actually do so for a large part of the night. Yet, dream recall shows large interindividual variability. Understanding the factors that influence dream recall is crucial for advancing our knowledge regarding dreams’ origin, significance, and functions. Here, we tackled this issue by prospectively collecting dream reports along with demographic information and psychometric, cognitive, actigraphic, and electroencephalographic measures in 204 healthy adults (18-70 y, 113 females). We found that attitude towards dreaming, proneness to mind wandering, and sleep patterns are associated with the probability of reporting a dream upon morning awakening. The likelihood of recalling dream content was predicted by age and vulnerability to interference. Moreover, dream recall appeared to be influenced by night-by-night changes in sleep patterns and showed seasonal fluctuations. Our results provide an account for previous observations regarding inter- and intra-individual variability in morning dream recall.

## Introduction

Dreams are subjective conscious experiences generated by the brain during sleep — when individuals are largely (though not completely; [1]) disconnected from the external environment on the sensory (input) and motor (output) sides and are typically unable to exert volition and self-reflection [2]. Dream experiences draw on previously acquired memories and beliefs and, thus, present relevant aspects of continuity with thoughts, concerns, and salient experiences of our waking self. In light of this, they are believed to represent an important window on —and to potentially have a direct role in— sleep-dependent processes involving learning and memory consolidation. Moreover, dreams have a tight relationship with psychophysical health. In fact, alterations in the frequency or content of oneiric experiences may accompany, or even precede, the waking manifestation of clinical symptoms related to psychiatric and neurological disorders [3], [4], [5]. Finally, the study of dreaming and dreamless sleep is regarded as a fundamental experimental model in the search for the functional bases of human consciousness. Indeed, as compared to task-based protocols exploring wakefulness conscious experiences, the study of dreams is naturally less influenced by confounding effects such as changes in attention, stimulus and task processing, task performance, and response preparation [6], [7].

In the 1950s, with the discovery of rapid eye movement (REM) sleep, researchers initially thought to have identified the neural correlates of dreaming [8], [9], as dream experiences appeared to be far more common in this stage than in non-REM (NREM) sleep. This idea fits well with the fast, low-amplitude electroencephalographic (EEG) activity similar to wakefulness that characterizes REM sleep, as opposed to the slow, high-amplitude activity of NREM sleep. However, later studies partially amended this view. Indeed, serial-awakening laboratory investigations determined that contentful dreams are reported on average following ∼85% of the awakenings from REM sleep and ∼45% of the awakenings from NREM sleep (e.g., [10]).

While the sleep stage preceding the awakening is considered a key determinant for whether or not a dream will be reported, evidence indicates that dream recall probability fluctuates greatly both within and across individuals [11]. Such a variability attracted public and scientific attention during the recent pandemic, when an abrupt surge in morning dream recall was reported worldwide [12]. Yet, our current understanding of the factors influencing dream generation and recall is scarce. Previous studies suggested that factors such as a positive attitude towards dreaming, frequent daydreaming, a high level of anxiety, female gender, and young age may be associated with higher dream recall frequency. Importantly, though, the available evidence is mostly based on retrospective measures potentially affected by biases such as memory- and personality-related distortions. Prospective studies conducted so far are sparse and hampered by significant limitations. Indeed, due to their higher costs, these investigations were typically performed on relatively small samples and explored only one or few variables potentially affecting dream recall.

This picture is further complicated by the inherent foundation of dream studies, to some degree, on the assumption that reports provided by individuals upon awakening are a reliable reflection of dream occurrence and content. However, any generated dream must be encoded in memory, and such a memory has to be later retrieved during wakefulness in order for a dream experience to be successfully recalled [11]. This issue is of particular importance given that memory processes appear to be altered during sleep and the subsequent period of *sleep inertia*. Indeed, individuals often wake up with the distinct feeling of having been dreaming but are unable to recall any detail of their experience. In some cases, the memory of the dream may be present at the moment of awakening but is rapidly lost if the experience is not immediately reported. These so-called ‘*white dreams*’ have been interpreted as reflecting a failure of memory encoding or retrieval [13], [14]. Yet, previous investigations provided little or no support for a direct relationship between memory skills and dream content recall [15], [16], [17], [18].

Here we set out to investigate the intra- and -inter-individual factors associated with morning dream recall in a large multimodal database collecting dream reports along with demographic information and psychometric, cognitive, actigraphic, and EEG measures. In this prospective study, a cohort of healthy adults recorded a report of their last dream experience each morning upon spontaneous awakening at home for 15 days (Fig. 1a). Sleep-wake patterns were tracked through actigraphy. A sub-sample also wore a portable EEG device at night. Moreover, all participants were characterized across a wide range of cognitive and psychological dimensions.

**Figure 1.**
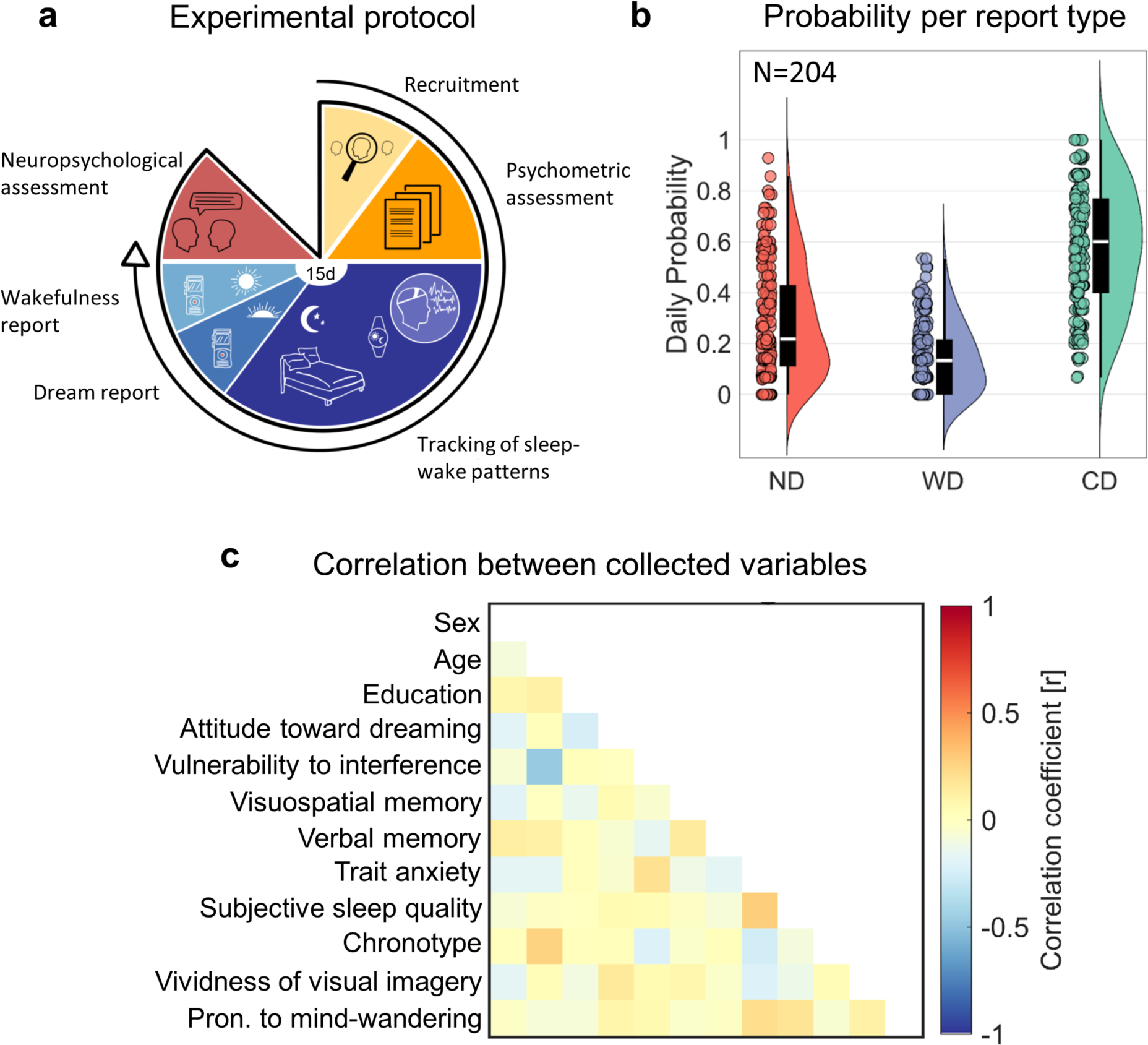
**a**, Outline of the experimental paradigm. **b**, Proportion of no dream experience (ND), white dream (WD), and contentful dream (CD) reports. For each report type, the corresponding raincloud plot (individual data points and probability distribution) and box plot are shown. On each box, the central mark indicates the median, and the bottom and top edges of the box indicate the 25th and 75th percentiles, respectively. The whiskers extend to the most extreme data points not considered outliers. **c**, Correlation (Spearman’s rho) between demographic, psychological and cognitive variables derived from questionnaires and tests.

## Results

A total of 204 participants sampled from the healthy Italian adult population (mean age 35.1 ± 12.5 y; 113 females, 55.4%) and 2900 morning reports were included in the analyses (14.22 ± 1.44 reports per subject). All participants wore an actigraph during the experimental period. Forty-two of them also wore a portable EEG device during their sleep (24 females; age 30.0 ± 5.2 y, range 22-44 y). Participants completed questionnaires and cognitive tests aimed at characterizing their attitude towards dreaming, trait anxiety levels, verbal memory, visual memory, vulnerability to interference, vividness of visual imagery, proneness to mind-wandering, subjective sleep quality, and subjective circadian preference.

### Frequency of morning dream reports

All morning verbal reports were evaluated and classified as either contentful dream experience (CD) if the verbal description included at least one reference to any kind of semantic content, dream experience without recall of content (‘*white dream*,’ WD) if the subject reported the perception of having dreamt but could not recall any feature of the experience, and no dream experience (ND) if the subject woke up with the feeling of not having dreamt. Moreover, the sum of dream experiences with and without recall of content (CD+WD) was computed to obtain an estimate of all cases where participants reported having dreamt. We assumed this metric to represent the best measurable estimate of generated dreams.

Fig. 1b shows the proportion of morning reports corresponding to CD (mean ± standard deviation = 0.58 ± 0.24), WD (0.14 ± 0.13), and ND (0.28 ± 0.22). On average, CD+WD probability was 0.72 ± 0.22, corresponding to 5.04 ± 1.54 dreams per week, in line with previous observations concerning dream recall frequency [19]. Morning dream recall frequency computed from the verbal diary was significantly higher than self-reported dream recall frequency (2.66 ± 2.29; signed-rank test, p < 0.0001; |g| = 1.19, CI = [1.04, 1.36]). The two indices of dream recall showed a moderate positive correlation (Spearman’s correlation, p < 0.0001; r = 0.46, CI = [0.35, 0.55]). These findings are consistent with previous studies indicating an incomplete correspondence between retrospective and prospective measures of dream recall [11].

### Inter-individual predictors

Next, we performed mixed-effect logistic regression analyses to identify potential predictors of dream generation (CD+WD vs. ND) and dream content memory (CD vs. WD). The following predictors were selected based on previous literature: age, sex, years of education, attitude towards dreaming, vulnerability to cognitive interference, verbal memory, visuospatial memory, trait anxiety, vividness of visual imagery, proneness to mind-wandering, self-reported sleep quality, and self-reported chronotype (Fig. 1c). Moreover, objective sleep measures were derived from actigraphic data. In particular, a principal component analysis (PCA) was performed across 24 distinct actigraphic indices (see Methods). We obtained four PCs together explaining 87.74% of the total variance (PC1 = 50.9%; PC2 = 15.4%; PC3 = 11.2%; PC4 = 10.4%; Fig. 2). To facilitate the interpretation of the PCs, we employed mixed-effect models including sleep-structure measures obtained in the subsample of participants who wore the portable EEG system during the experimental nights (Tables S2-5). We found that PC1 was positively associated with the proportion of wakefulness (q = 0.001, False Discovery Rate -FDR-correction) and N1 (q = 0.001), as well as negatively associated with the proportion of N2 and N3 sleep (q =0.016; model adjusted R^2^ = 0.47). Moreover, PC2 was negatively associated with the proportion of N3 sleep (q < 0.001) and W (q = 0.042; model adjusted R^2^ = 0.41), whereas PC4 was negatively associated with the proportions of N2, N3, and REM sleep (q < 0.005; model adjusted R^2^ = 0.38). No significant predictors were identified for PC3. Based on these observations (Fig. 2b) and the distribution of PC loadings (Fig. 2a), the four PCs could be assumed to mainly reflect, respectively, *sleep fragmentation* (PC1), *long, non-N3 sleep* (PC2; hereinafter indicated as ‘*long light sleep*’), *stable sleep with advanced phase* (PC3; ‘*stable advanced sleep*’), and *unstable sleep with advanced phase* (PC4; ‘*unstable advanced sleep*’).

**Figure 2.**
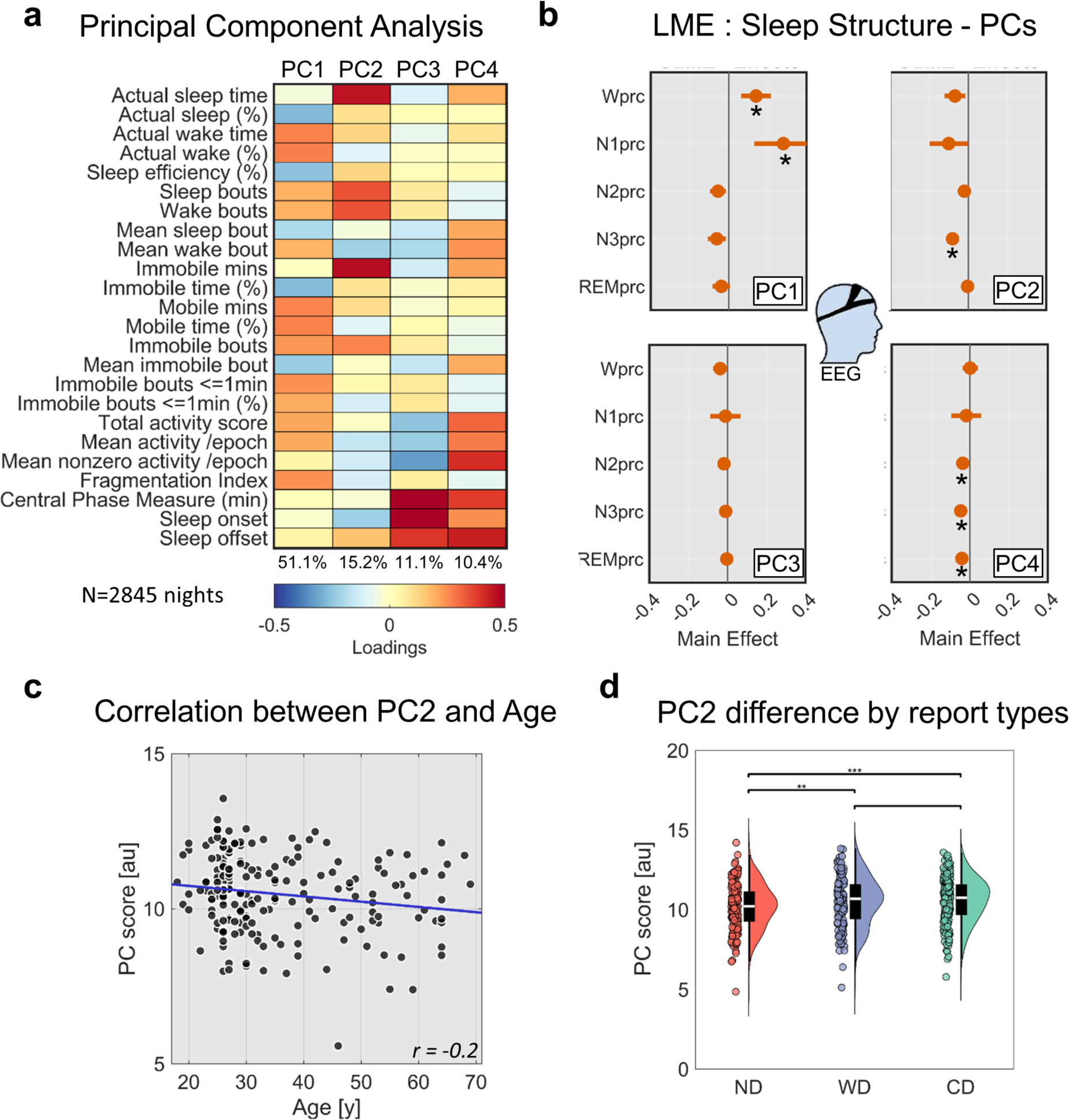
**a**, Principal Component (PC) analysis of actigraphic data (N = 2845 nights). We identified four PCs explaining at least 10% of the variance. For each PC, the plot shows the loadings on each actigraphic metric and the explained variance (bottom). **b**, Linear Mixed Effect (LME) models were applied to investigate the EEG-derived sleep-structure parameters associated with each PC (* marks significant effect at q < 0.05, FDR correction; N = 483 nights). Age and Sex were accounted for in the models, but their effects are not shown. **c**, Correlation between PC2 loadings and age. A significant negative correlation was observed. **d**, Comparison of mean PC2 loadings associated with nights followed by no dream experience (ND), white dream (WD), and contentful dream (CD) reports (* p < 0.05; ** p < 0.01; *** p < 0.001).

We found that morning dream reports were predicted by attitude towards dreaming, proneness to mind wandering, and long light sleep (q < 0.05, FDR correction; adjusted R^2^ = 0.17; Fig. 3, also see Table S6). Contrary to previous research [19], [20], we did not find significant effects of age and sex on dream recall (i.e., higher recall in younger individuals and females). However, we noted a significant relationship between sex and attitude towards dreaming, with the latter being higher in females (N=113, 36.6 ± 13.4 y) compared to males (N=91, 33.3 ± 11.2 y; rank-sum test, p = 0.014; |g| = 0.34, CI = [0.06, 0.62]). Moreover, we found a significant negative correlation between light sleep (PC2) and age (Spearman’s correlation, p = 0.008; r = −0.19, CI = [−0.32, −0.06]). These results suggest that previously described effects of age and sex could have been mediated by other factors. The distinction between CD and WD was instead predicted by age and vulnerability to interference (q < 0.05; adjusted R^2^ = 0.11; Fig. 3b, also see Table S7).

**Figure 3.**
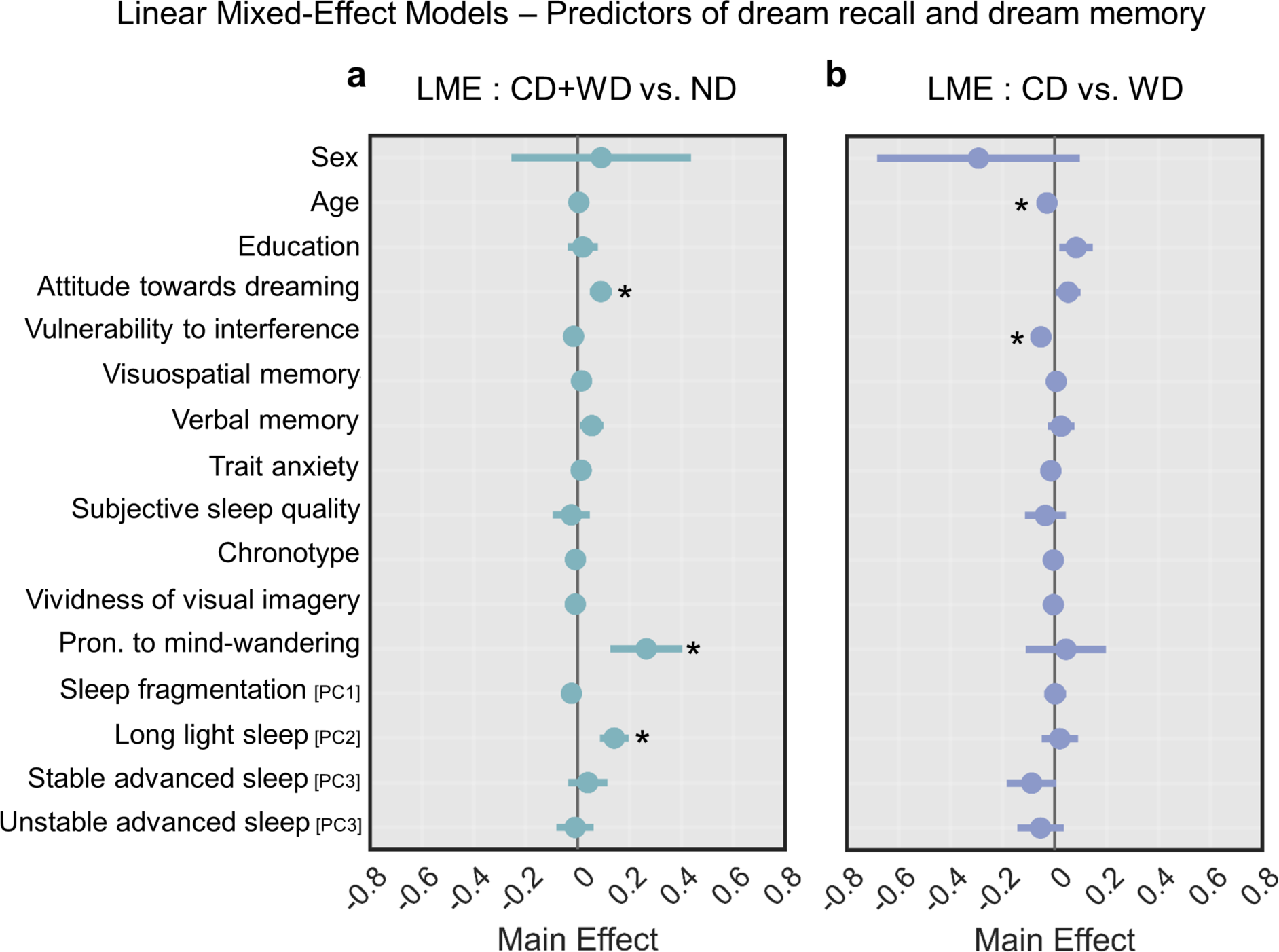
Linear Mixed Effect models exploring the inter-individuals predictors of morning dream recall (**a**; N = 2900) and dream content memory (**b**; N = 2077). The overall effects are shown for each variable included in the models. * mark significant effects at q < 0.05, FDR correction.

### Night-by-night variations

Next, we investigated how sleep patterns affect night-by-night variations in morning dream recall. We thus compared mean actigraphic PCs for nights followed (CD+WD) or not (ND) by a morning dream report. No significant differences were found for sleep fragmentation (PC1; p = 0.102, uncorrected; N = 182), stable advanced sleep (PC3; p = 0.992), and unstable advanced sleep (PC4; p = 0.969). Instead, long light sleep (PC2) scores were significantly higher for CD+WD relative to ND (N = 182; corrected q < 0.001; |g| = 0.28 -[0.17, 0.42]). Therefore, we further investigated for this PC potential differences among CD, WD, and ND. We found that both WD (N = 129; q = 0.006; |g| = 0.22, CI = [0.07, 0.38]) and CD (N = 182; q < 0.001; |g| = 0.26, CI = [0.14, 0.38]) had significantly higher scores relative to ND, whereas no differences were found between WD and CD (N = 140; p = 0.806, uncorrected). Overall, these results indicate that CD and WD depend on similar sleep patterns and suggest that long light sleep may be associated with dreaming per se, rather than dream content memory.

In addition, we investigated the impact of sleep structure measures on dream recall in the sample of participants who wore the portable EEG system. We found that morning dream recall tended to be associated with the proportion of overnight REM sleep (N = 37; uncorrected p = 0.027), but this effect did not survive correction for multiple comparisons (q = 0.135; |g| = 0.38, CI = [0.05, 0.77]). A follow-up contrast among CD, WD, and ND reports showed a significant difference in the amount of REM sleep between CD and ND (N = 37; uncorrected p = 0.023; corrected q = 0.070; |g| = 0.39 - [0.08, 0.77]) but not between WD and ND (N = 25; p = 0.276) or between CD and WD (N = 28; p = 0.466). It is important to note that this analysis concerned the sleep macrostructure of the entire night and that a longer REM duration does not necessarily imply that participants woke up from this stage. However, similar analyses performed using the last 2 hours of sleep yielded similar results (CD+WD vs. ND, p = 0.017; CD vs. ND, p = 0.007; CD vs. WD, p = 0.029; WD vs. ND, p = 0.696).

Overall, the obtained results suggest that individuals may be more likely to recall dreams when they wake up from long sleep nights with a small proportion of deep, N3 sleep and higher REM content. This finding is consistent with previous observations indicating a negative correlation between sleep stages with a high slow wave activity (N3) and dreaming [17], [21].

### Seasonal variations

Owing to the fact that data collection took place over a period of four years (from 2020 to 2024), we investigated potential fluctuations in morning dream recall across seasonal cycles (Fig. 4). For this analysis, morning dream report (CD+WD) rates were computed for each participant and adjusted for age, attitude towards dreaming, proneness to mind wandering and mean light sleep (PC2) scores. We found that morning dream report probability was significantly lower in Winter relative to Spring (uncorrected p = 0.005, corrected q = 0.019; |g| = 0.59, CI = [0.19, 1.04]). A trend towards a lower morning dream recall in Winter relative to Autumn was also observed (uncorrected p = 0.037, corrected q = 0.074; |g| = 0.44, CI = [0.10, 0.84]).

**Figure 4.**
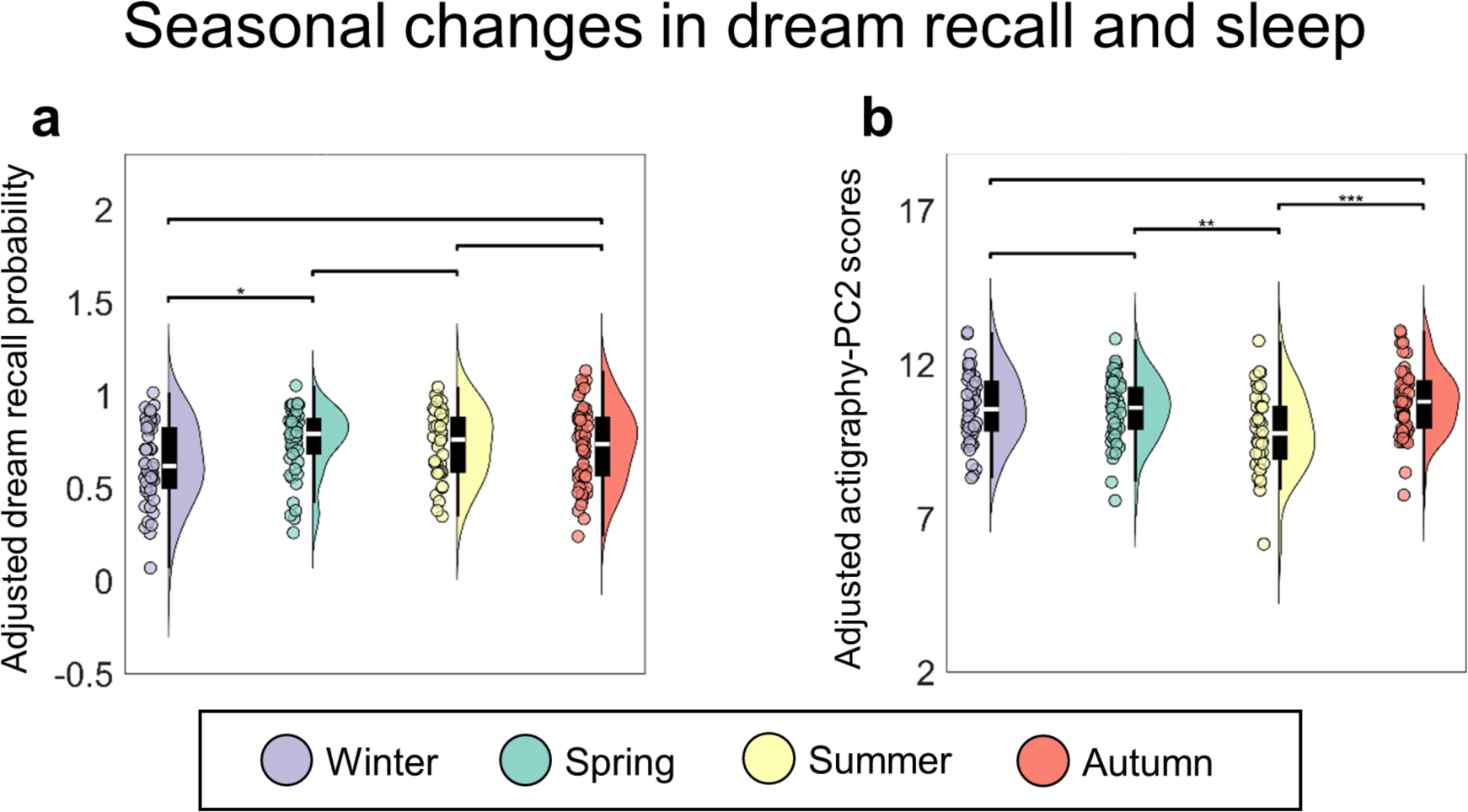
Changes in morning dream recall (**a**) and sleep patterns reflected by the actigraphy-PC2 (**b**) across seasons. * q < 0.05, ** q < 0.01, *** q < 0.001 (FDR correction). Dream recall probability values were adjusted for age, sex, vulnerability to interference, attitude towards dreaming, and PC2. PC scores were adjusted for participants’ age and sex.

To determine whether the observed seasonal changes could mirror changes in sleep patterns as assessed using actigraphic indices, we further analyzed potential seasonal variations in PC scores. For these analyses, mean PC scores were adjusted for age and sex. We found no significant seasonal changes for sleep fragmentation (PC1; uncorrected p > 0.074), stable advanced sleep (PC3; p > 0.175), and unstable advanced sleep (PC4; p > 0.195). A significant impact of the season was instead found for long light sleep (PC2). In particular, long light sleep scores were lower in Summer relative to both Spring (uncorrected p = 0.005, corrected q = 0.005; |g| = 0.60, CI = [0.19, 1.07]) and Autumn (uncorrected p < 0.001, corrected q < 0.001; |g| = 0.81, CI = [0.42, 1.27]). Overall, seasonal sleep changes did not appear to mirror relative changes in dream recall across seasons.

## Discussion

Here, we show that the likelihood of waking up in the morning from a subjective dream experience is predicted by three main factors, that are attitude towards dreaming, proneness to mind wandering, and trait differences in overnight sleep patterns. Moreover, individual differences in the tendency to recall the content of dream experiences as opposed to the mere awareness of having dreamt are predicted by vulnerability to interference and age.

A positive association between attitude towards dreaming and dream recall has been consistently described both by studies employing retrospective measures, such as questionnaires, and by prospective investigations performed, among other approaches, by means of dream diaries [22]. Yet, the causal relationship still represents an open question. Indeed, it has been suggested that a pre-existing interest in dreams may drive a person to apply strategies aimed at increasing successful dream retrieval (e.g., keeping a dream diary). On the other hand, individuals who often remember their dreams may develop an interest in their possible meaning or significance. Notably, our findings indicate that while attitude towards dreams influences the likelihood of reporting the experience of a dream, it does not significantly impact the probability of recalling dream content. This observation lends indirect support to the notion that the association between attitude towards dreaming and dream recall may be driven by factors beyond mere memory processes.

The tendency towards mind wandering emerges in our study as another robust positive predictor of dream recall. A relevant perspective posits that dreaming and mind-wandering, or daydreaming, may exist along a continuum, relying on similar brain functional mechanisms and structures [23], [24], [25], [26]. Recent research has highlighted the involvement of overlapping neural networks, particularly the default mode network (DMN), in both mind wandering and dreaming [24], [27]. The DMN, including brain regions such as the medial prefrontal cortex and posterior cingulate cortex, is known to be active during periods of internally focused cognition and self-referential thought [28]. Given its role in introspective mental processes, the DMN has been implicated in promoting mind wandering during wakefulness [29], [30]. Consequently, the association between mind wandering and dream recall observed in our findings may indicate a heightened propensity to spontaneously generate dream-like experiences, irrespective of external stimuli and vigilance states. A possible alternative interpretation is that individuals who engage in more frequent daydreaming may pay greater attention to their internal states and subjective experiences. Such a heightened introspective awareness might in turn facilitate the encoding and recall of dream experiences.

Our analyses revealed an important role of overnight sleep patterns as extracted from actigraphic data. Specifically, we found that individuals who typically have long light sleep episodes coupled with a low proportion of deep N3 sleep, exhibit a heightened probability of dream recall compared to those experiencing shorter, N3-rich sleep. This observation aligns with prior research describing a negative relationship between the occurrence of slow waves typical of NREM sleep and dreaming [31]. Indeed, not only the probability of reporting a dream upon awakening decreases in parallel with the increase in slow wave activity during the deepening of NREM sleep, from N1 to N3 [10], but, even within the same sleep stage (be that NREM or REM), higher slow wave activity is more often observed when individuals report no dream experiences [31], [32], [33], [34]. Sleep slow waves are primarily local events that result from an oscillation of cortical neuronal populations between a depolarized, active state, and a hyperpolarized, silent state. Their asynchronous occurrence across cortical areas is thought to cause a breakdown in cortical connectivity and impair information integration, a key prerequisite for the emergence of consciousness [35], [36]. Importantly, slow waves are homeostatically regulated, so that they increase in number and amplitude after sleep deprivation/restriction and decrease across sleep cycles, during a night of sleep (or a nap; [37]). In this light, a large amount of N3 sleep may indicate a high sleep pressure, with a strong slow wave activity that may decrease the probability of experiencing dreams regardless of the sleep stage from which the sleeper wakes up.

In addition, to the best of our knowledge, this is the first study to demonstrate that dream recall frequency is characterized by seasonal fluctuations, being lower during Winter as compared to Spring and Autumn. While our actigraphic data did not allow us to detect macrostructural sleep changes that could explain the variations in dream recall, we were not able to investigate possible finer variations in sleep macro- or micro-structure. Such variations could contribute to determining the decrease in dream recall observed in Winter and should be the object of further investigations.

Next, we investigated the individual factors that may affect the retrieval of dream content and lead to the perception of having been dreaming without managing to recall any feature of that experience, so-called *white dreams* [12]. Notably, we observed that the same sleep patterns associated with morning dream recall -namely, long, light sleep bouts-are equally associated with recalling dream content and forgetting it. This observation suggests that white dreams and contentful dreams both reflect true, generated dreams with different degrees of recall of specific aspects of the experience. Interestingly, in line with previous work, we did not find potential associations between white dreams and visual or verbal memory [13], suggesting that memory processes regarding dream content may not be affected by general memory skills. Instead, we found that individuals with a higher vulnerability to interference tend to more often forget the content of their dreams upon awakening. This observation is consistent with previous evidence suggesting that the memory of a dream may be lost if interference occurs between dream experience and retrieval [38]. Indeed, higher resilience to interference may allow individuals to maintain the focus of attention and memory on the dream in spite of situational (e.g., turning off the alarm sound or talking with the bed partner) or internal (e.g., thoughts about the upcoming schedule or thinking about current concerns) interferences.

Previous work suggested that dream recall may decrease with age [19], [39] and that women may recall more dreams than men [19], [20], [40]. Neither of these findings was confirmed here. We hypothesize that such effects might be actually mediated by other variables. Indeed, for instance, we found that the attitude towards dreaming, positively associated with dream recall, is higher in women than in men [41]. Moreover, our results suggest that aging may be associated with changes in sleep patterns —and in particular with a decrease in the long, light sleep bouts— that may in turn affect dream generation processes. However, we found an independent effect of age on content recall, so that aging is associated with a higher probability of reporting *white dreams*. The mechanism underlying this association is unclear, and such an effect may actually be mediated by variations in other cognitive processes not investigated here, such as working memory skills (but see [42]).

In conclusion, our study demonstrates that specific inter-subject (trait) and intra-subject (state) variables influence the likelihood of having and recalling a dream experience. Notably, our findings show that similar overnight sleep patterns increase the probability of both contentful and *white dreams*, and that the memory retention for dream content may be primarily lost due to interference by external or internal factors. These observations support the notion that *white dreams* represent actual dream experiences, with memories of their content fading upon waking.

## Materials and Methods

### Participants

The study was conducted on a sample of 217 healthy Italian native language speakers from 18 to 70 years old (116 females). Of these, ten failed to comply with the experimental protocol, and three provided less than seven recordings (see below) leading to a final sample of 204 participants. Data collection was carried out between March 2020 and March 2024, covering a period of four years. Only individuals with regular sleep/wake patterns, six to eight hours of sleep per night, and no diagnosis of sleep-related problems or of any other pathological condition that might have compromised their sleep were recruited in the study. Moreover, we excluded volunteers who were taking medications that could have affected sleep patterns at the time of the study and individuals who had a recent (last 6 months) history of alcohol and drug abuse. Finally, women who were pregnant, were planning a pregnancy, or were breastfeeding at the time of the study were also excluded.

Each study participant went through three phases (see below): i) a screening interview followed by the completion of a questionnaire battery, ii) an experimental stage where sleep patterns and morning reports of subjective sleep conscious experiences were collected for 15 days, and iii) a final session with the administration of a battery of cognitive tests.

The study was conducted under a protocol approved by the Local Joint Ethical Committee for Research (#11/2020). All volunteers signed a written informed consent form before taking part in the study and retained the faculty to drop from the study at any time.

### Screening interview and self-assessment questionnaires

All volunteers underwent an anamnestic interview aimed at assessing their general health and adherence to inclusion/exclusion criteria. Recruited participants were then asked to fill out several questionnaires aimed at investigating their attitude towards dreaming [41], trait anxiety levels (*State-Trait Anxiety Inventory, STAI*; [43]), vividness of visual imagery (*Vividness of Visual Imagery Questionnaire, VVIQ*; [44]), proneness to mind-wandering (*Mind Wandering - Spontaneous and Deliberate Scale, MW*; [45]), subjective sleep quality (*Pittsburgh Sleep Quality Index, PSQI*; [46]), subjective circadian preference (*Morningness-Eveningness Questionnaire*; [47]). Attitude towards dreaming was assessed using a 6-item questionnaire where participants were asked to provide their degree of agreement with six statements regarding the general meaning and significance of dreams on a Likert Scale from 0 (‘completely disagree’) to 4 (‘completely agree’). Three items were positive statements about dreams (e.g. ‘dreams are a good way of learning about my true feelings’) and three were negative (e.g. ‘dreams are random nonsense from the brain’). A global score was computed by subtracting the sum of scores provided to the negative statements from the sum of scores associated with the positive statements. Finally, participants completed a questionnaire about their dream experiences in the previous three months, which included one item aimed at assessing the frequency of morning dream recall [48].

### Collection of morning dream reports and sleep patterns

Volunteers who met all the inclusion criteria were provided with an actigraph and a voice-recorder and were asked to record each morning, upon awakening from sleep, everything that was going through their mind just before they woke up, everything they remembered, every experience or thought they had before awakening. It is important to note that this procedure partially differs from those commonly used in home-based experiments on dreams, in which participants are asked to report all dreams that they had during a given sleep night. Indeed, we specifically chose to focus on the very last experience the subjects had before awakening in order to minimize the impact of confounding effects that may intervene between the dream experience and the report. Furthermore, at pseudo-random times during the day, participants were also contacted and asked to record everything that was going through their minds up to 15 minutes before they started the recording. In particular, a simple phone text message containing the word “record” (“*registra*”) was sent to the volunteers at pseudo-random times during the day (Fig. 1a). Wakefulness reports were not analyzed in the present study.

Only subjects who provided at least seven reports during the experimental period were included in our analyses. This selection criterion led to the exclusion of three participants. One participant who reported altered sleep-wake patterns (sleep restriction) in three nights voluntarily extended the experimental period to 20 days. We discarded the reports provided by this participant at the awakening from the altered nights and on the experimental days immediately following. Furthermore, we excluded delayed dream recall reports. Though subjects were specifically required to only report the experiences they remembered right after the awakening, occasionally they retrieved and recorded the dream experience later during the day. Since these memories might be triggered and distorted by external stimuli and events experienced during wakefulness, we chose to exclude those data.

During the 15 days of the study, participants wore an actigraph to track sleep-wake patterns (MotionWatch-8, Camtech). A subgroup of 50 volunteers (27 females; age 29.7 ± 5.2 y, range 22-44 y) also had their sleep-related brain activity recorded through a portable EEG system (*DREEM*) equipped with five EEG dry electrodes (seven derivations: Fp1-O1, Fp1-O2, Fp1-F7, F8-F7, F7-O1, F8-O2, Fp1-F8), a pulse sensor, and a 3D accelerometer. Eight participants interrupted EEG data collection due to discomfort while sleeping. Therefore, we were able to analyze data collected from 42 participants.

### Cognitive testing

At the end of the two-week period, all participants underwent a neuropsychological assessment aimed at evaluating different cognitive abilities. The neuropsychological tests comprised: *Stroop Color and Word Test*, for assessing participants’ processing speed and vulnerability to cognitive interference (SCWT; [49]); *Babcock Story Recall Test* (immediate recalling – delayed recalling), for evaluating participants’ episodic and verbal memory (*BSRC*; [50]); *Rey–Osterrieth complex figure* (immediate copy - delayed copy), for evaluating participants’ visuo-constructional ability and visual memory (ROCF; [51]).

### Analysis of actigraphic data

Actigraphic recordings were evaluated by means of the *MotionWare* Software [52]. The following 22 measures were computed (as described within the software user guide): *actual sleep (or wake) time* (the total time spent in sleep/wake according to the epoch-by-epoch wake/sleep categorisation); *actual sleep (or wake) percent* (actual sleep/wake time expressed as a percentage of the total elapsed time between ‘*fell asleep*’ and ‘*wake up*’ times); *sleep efficiency* (actual sleep time expressed as a percentage of time in bed); *sleep (or wake) bouts* (the number of contiguous sections categorized as sleep/wake in the epoch-by-epoch wake/sleep categorisation); *mean sleep (or wake) bout* (the average length of each of the sleep/wake bouts); *immobile (or mobile) minutes* (the total time categorized as immobile/mobile in the epoch-by-epoch mobile/immobile categorisation); *percentage of immobile (or mobile) time* (the immobile/mobile time expressed as a percentage of the assumed sleep time); *immobile bouts* (the number of contiguous sections categorized as immobile in the epoch-by-epoch mobile and immobile categorisation); *mean immobile bout* (the average length of each of the immobile bouts); *immobile bouts <=1min* (the number of immobile bouts which were less than or equal to one minute in length); *percentage of immobile bouts <=1min* (the number of immobile bouts less than or equal to one minute expressed as a percentage of the total number of immobile bouts); *total activity score* (the total of all the activity counts during the assumed sleep period); *mean activity/epoch* (the total activity score divided by the number of epochs in the assumed sleep period); *mean nonzero activity per epoch* (the total activity score divided by the number of epochs with greater than zero activity in the assumed sleep period); *fragmentation index* (the sum of the ‘*percentage of mobile time*’ and the ‘*percentage of immobile bouts <=1min*’); *central phase measure* (the midpoint between the ‘*fell asleep*’ and ‘*wake up*’ times, expressed as the number of minutes past midnight). Moreover, we expressed the ‘*fell asleep*’ and the ‘*wake up*’ *times* as the number of minutes past midnight.

We discarded measures from single nights where the actigraph appeared to have been removed (e.g., cases where participants removed it and forgot to wear it again before sleep time). In particular, we discarded nights that met at least three of the following heuristic criteria: number of ‘*immobile bouts <=1min*’ ≤ 2, number of ‘*sleep bouts*’ ≤ 4, ‘*fragmentation index*’ ≤ 1.5%, ‘*actual sleep percent*’ ≥ 96 %. Overall, we excluded actigraphic recordings collected in 39 nights across 28 participants due to missing or unreliable data. Moreover, actigraphic data (all nights) was lost in four participants due to technical issues (4 females, age 22-30 y).

Given that the actigraphic variables were highly correlated with each other, we applied dimensionality reduction through principal component analysis (PCA; N = 2845 nights). We retained PCs that explained at least 10% of the total variance.

### Analysis of portable EEG system data

Data collected using the *DREEM* device were analyzed using the associated automated sleep scoring software [53]. The obtained hypnograms were used to compute the percentages of wakefulness, N1, N2, N3, and REM sleep for each night (N = 483). Then, sleep structure measures were used to facilitate the interpretation of actigraphy-related PCs. Specifically, we employed mixed-effect models to explore the association between sleep structure measures and each of the four PCs. Four identical, independent models were used. An FDR correction [54] for multiple comparisons was applied to adjust p-values assigned to each of the tested predictors.

Below we reported the adopted model d using Wilkinson’s notation, where Y represents the predicted variable (i.e., each PC), Subj is the participant identification number and Night is the experimental night counted from the beginning of the experiment:

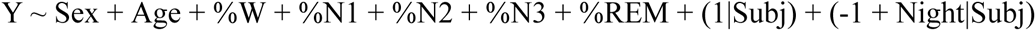

### Predictors of dream recall and dream content memory

Two separate mixed-effect logistic models were used to investigate the inter-individual predictors of dream recall (CD+WD vs. NE) and dream content memory (CD vs. WD). The same predictors were included in the two models: age, sex, education, attitude towards dreaming (ATD), trait anxiety (STAI), vulnerability to interference (SCWT), vividness of visual imagery (VVIQ), proneness to mind-wandering (MW), verbal memory (BSRT), visual memory (ROCFr), subjective sleep quality (PSQI), subjective circadian preference (MEQ), and four actigraphy-derived PCs (see Results). The models also accounted for possible effects of the experimental days when the reports were collected.

The first model, aimed at predicting morning dream recall (CD+WD) as compared to ND reports, included a total of 2900 reports across 204 participants. The second model, aimed at predicting contentful dream recall (CD) as compared to WD reports, included a total of 2077 reports across 204 participants. An FDR correction [54] for multiple comparisons was applied to adjust p-values assigned to each of the tested predictors.

Similarly to the analysis of EEG data, Subj and Night variables were included as random-effect terms. Below is the model used:

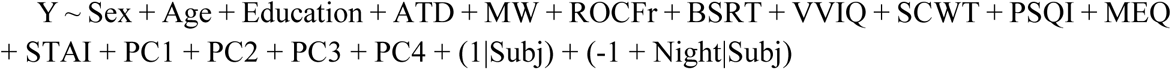

### Night-by-night variations in dream recall

We investigated how sleep patterns affect night-by-night variations in morning dream recall. Similar analyses were performed using actigraphy-derived PC-scores and sleep structure measures obtained from EEG data. Specifically, we first averaged values for nights associated with a dream report (CD+WD) and nights that were not followed by a dream report (ND). Then mean values were compared across report types using non-parametric sign-rank tests for paired samples. In case a significant effect was found for a specific PC or sleep structure parameter, additional comparisons were carried out across CD, WD, and NE report types. FDR corrections were applied to account for multiple comparisons.

### Seasonal variations in dream recall

To investigate the impact of seasonal variations on morning dream recall, we estimated morning dream recall probability per subject and adjusted the obtained values for age, sex, attitude towards dreaming, proneness to mind-wandering, and long light sleep (PC2) scores (see Results). Then, we grouped subjects according to the season (Winter, Spring, Summer, Autumn) in which they carried out the study, using as a reference the central day of their experimental period. Rank-sum tests were performed to compare adjacent seasons and an FDR correction was applied to account for multiple comparisons.

To determine whether the seasonal changes in dream recall could be explained by changes in sleep patterns, we analyzed potential seasonal variations in actigraphy-derived PC scores. For these analyses, mean PC scores were independently adjusted for age and sex. Possible effects were assessed as described above.

## Acknowledgments

This work was supported by a grant from the BIAL Foundation (#091/2020; to G.Be., M.B., and V.E.). The authors thank Elena Capriglia, Monica Di Giuliano, Francesco Lomi, and Aurora Salina for their help with data collection and preprocessing.

## Author Contributions

Conceptualization: G.Be., M.B., V.E.; Investigation: V.E., G.Bo., B.P.; Methodology: G.H., G.Be., V.E.; Software: G.H., G.Be., V.E., D.B.; Formal Analysis: V.E., D.B., G.Be., G.H.; Visualization: V.E., G.B.; Supervision: G.H., G.B.; Funding acquisition: G.Be., M.B.; Project administration: G.Be; Data Curation: V.E.; Writing - Original Draft: V.E., G.Be.; Writing - Review and Editing: All authors. All authors have read and agreed to the published version of the manuscript.

## Supplementary Materials

**Table S1.**
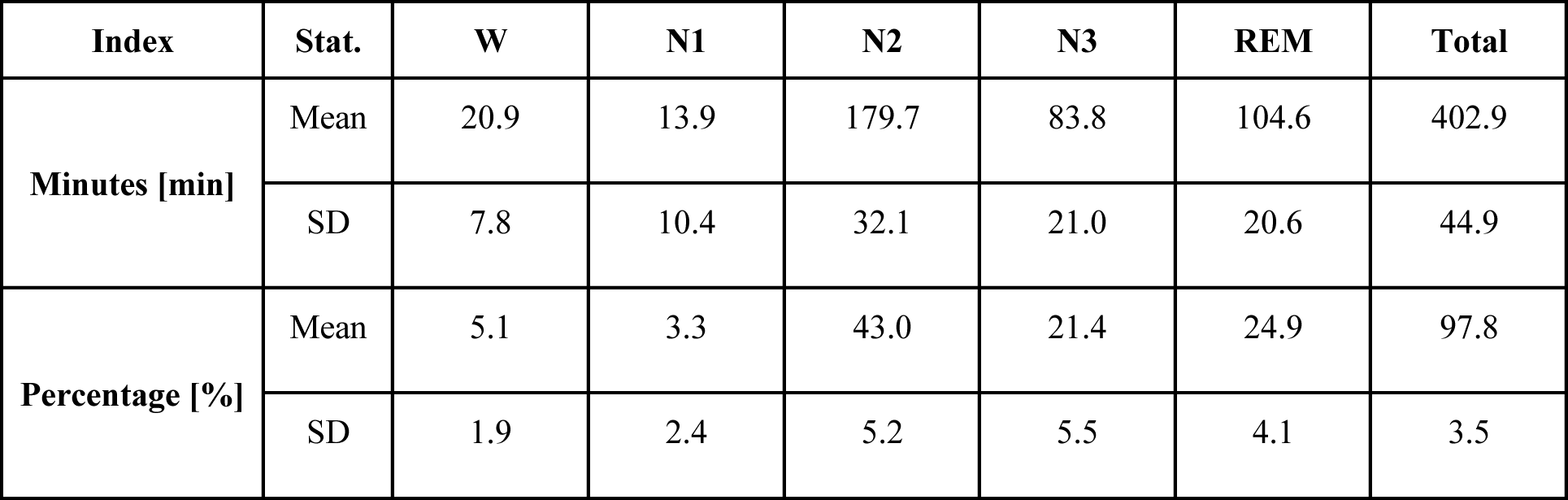
Sleep structure metrics computed for the 42 participants who wore the portable EEG device at night.

| Index | Stat. | W | N1 | N2 | N3 | REM | Total |
| --- | --- | --- | --- | --- | --- | --- | --- |
| Minutes [min] | Mean | 20.9 | 13.9 | 179.7 | 83.8 | 104.6 | 402.9 |
|  | SD | 7.8 | 10.4 | 32.1 | 21.0 | 20.6 | 44.9 |
| Percentage [%] | Mean | 5.1 | 3.3 | 43.0 | 21.4 | 24.9 | 97.8 |
|  | SD | 1.9 | 2.4 | 5.2 | 5.5 | 4.1 | 3.5 |

**Table S2.** LME results for PC1 (df = 450). For each predictor, we reported the estimate, standard error (SE), t statistics, p-value, and lower and upper bounds of the effect 95% confidence interval.

| **Name** | **Estimate** | **SE** | **tStat** | **pValue** | **Lower CI** | **Upper CI** |
| --- | --- | --- | --- | --- | --- | --- |
| **Intercept** | 0.034068 | 2.4362 | 0.013984 | 0.98885 | -4.7537 | 4.8218 |
| **Sex** | 0.9454 | 0.52002 | 1.818 | 0.009721 | -0.076567 | 1.9674 |
| **Age** | -0.042777 | 0.050732 | -0.94221 | 0.39936 | -0.14248 | 0.056923 |
| **Wprc** | 0.14052 | 0.0388 | 3.6217 | 0.000326 | 0.06427 | 0.21677 |
| **N1prc** | 0.28084 | 0.076199 | 3.6855 | 0.00025603 | 0.13109 | 0.43059 |
| **N2prc** | -0.05491 | 0.021017 | -2.677 | 0.0092843 | -0.096213 | -0.013607 |
| **N3prc** | -0.061246 | 0.023343 | -2.6237 | 0.0089934 | -0.10712 | -0.015371 |
| **REMPrc** | -0.037996 | 0.022757 | -1.6697 | 0.095676 | -0.082719 | 0.0067259 |

**Table S3.** LME results for PC2 (df = 450). For each predictor, we reported the estimate, standard error (SE), t statistics, p-value, and lower and upper bounds of the effect 95% confidence interval.

| **Name** | **Estimate** | **SE** | **tStat** | **pValue** | **Lower CI** | **Upper CI** |
| --- | --- | --- | --- | --- | --- | --- |
| **Intercept** | 16.064 | 1.6154 | 9.9439 | 3.4147e-21 | 12.889 | 19.238 |
| **Sex** | -0.013643 | 0.31516 | -0.04329 | 0.96549 | -0.63301 | 0.60572 |
| **Age** | -0.06604 | 0.030786 | -2.1451 | 0.032479 | -0.12654 | -0.005537 |
| **Wprc** | -0.068185 | 0.02706 | -2.5198 | 0.012088 | -0.12136 | -0.015005 |
| **N1prc** | -0.10027 | 0.049787 | -2.0141 | 0.044597 | -0.19812 | -0.0024304 |
| **N2prc** | -0.019915 | 0.014718 | -1.3531 | 0.1767 | -0.048839 | 0.0090096 |
| **N3prc** | -0.08055 | 0.016288 | -4.9454 | 1.075e-06 | -0.11256 | -0.04854 |
| **REMprc** | -0.0033869 | 0.01591 | -0.21288 | 0.83152 | -0.034654 | 0.02788 |

**Table S4.** LME results for PC3 (df = 450). For each predictor, we reported the estimate, standard error (SE), t statistics, p-value, and lower and upper bounds of the effect 95% confidence interval.

| **Name** | **Estimate** | **SE** | **tStat** | **pValue** | **Lower CI** | **Upper CI** |
| --- | --- | --- | --- | --- | --- | --- |
| **Intercept** | 3.582 | 1.3024 | 2.7503 | 0.0061942 | 1.0224 | 6.1415 |
| **Sex** | 0.2691 | 0.31692 | 0.8491 | 0.39628 | -0.35373 | 0.89193 |
| **Age** | -0.075666 | 0.030853 | -2.4525 | 0.014567 | -0.1363 | -0.015032 |
| **Wprc** | -0.037146 | 0.018587 | -1.9985 | 0.04626 | -0.073674 | -0.00061862 |
| **N1prc** | -0.010461 | 0.039865 | -0.26241 | 0.79312 | -0.088805 | 0.067883 |
| **N2prc** | -0.018081 | 0.010013 | -1.8057 | 0.071634 | -0.03776 | 0.0015976 |
| **N3prc** | -0.0092717 | 0.011166 | -0.83035 | 0.40678 | -0.031215 | 0.012672 |
| **REMPrc** | -0.0036555 | 0.010861 | -0.33658 | 0.73659 | -0.024999 | 0.017688 |

**Table S5.**
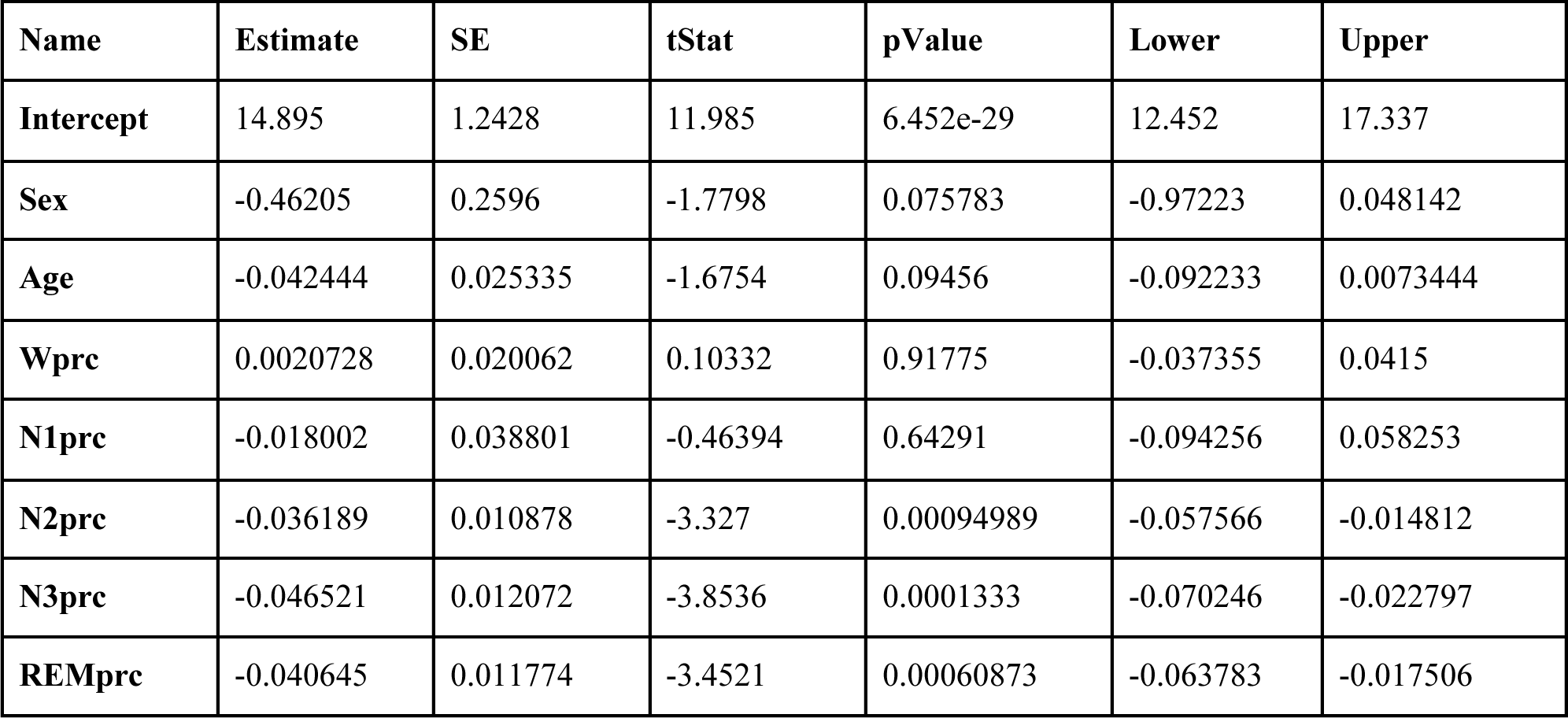
LME results for PC4 (df = 450). For each predictor, we reported the estimate, standard error (SE), t statistics, p-value, and lower and upper bounds of the effect 95% confidence interval.

**Table S6.**
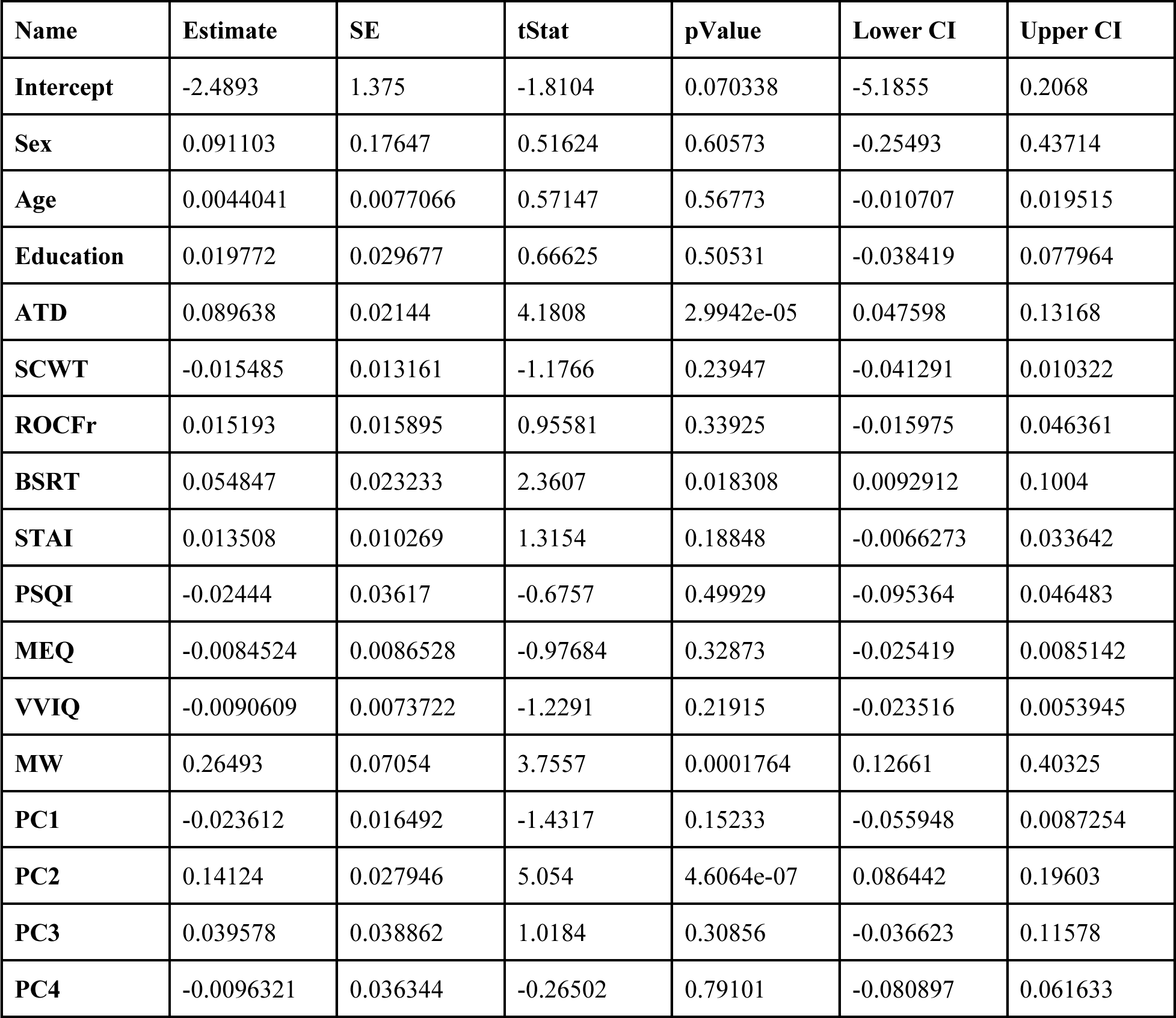
LME results for CD+WD vs. ND (df = 2776). For each predictor, we reported the estimate, standard error (SE), t statistics, p-value, and lower and upper bounds of the effect 95% confidence interval.

| Name | Estimate | SE | tStat | pValue | Lower CI | Upper CI |
| --- | --- | --- | --- | --- | --- | --- |
| **Intercept** | -2.4893 | 1.375 | -1.8104 | 0.070338 | -5.1855 | 0.2068 |
| **Sex** | 0.091103 | 0.17647 | 0.51624 | 0.60573 | -0.25493 | 0.43714 |
| **Age** | 0.0044041 | 0.0077066 | 0.57147 | 0.56773 | -0.010707 | 0.019515 |
| **Education** | 0.019772 | 0.029677 | 0.66625 | 0.50531 | -0.038419 | 0.077964 |
| **ATD** | 0.089638 | 0.02144 | 4.1808 | 2.9942e-05 | 0.047598 | 0.13168 |
| **SCWT** | -0.015485 | 0.013161 | -1.1766 | 0.23947 | -0.041291 | 0.010322 |
| **ROCFr** | 0.015193 | 0.015895 | 0.95581 | 0.33925 | -0.015975 | 0.046361 |
| **BSRT** | 0.054847 | 0.023233 | 2.3607 | 0.018308 | 0.0092912 | 0.1004 |
| **STAI** | 0.013508 | 0.010269 | 1.3154 | 0.18848 | -0.0066273 | 0.033642 |
| **PSQI** | -0.02444 | 0.03617 | -0.6757 | 0.49929 | -0.095364 | 0.046483 |
| **MEQ** | -0.0084524 | 0.0086528 | -0.97684 | 0.32873 | -0.025419 | 0.0085142 |
| **VVIQ** | -0.0090609 | 0.0073722 | -1.2291 | 0.21915 | -0.023516 | 0.0053945 |
| **MW** | 0.26493 | 0.07054 | 3.7557 | 0.0001764 | 0.12661 | 0.40325 |
| **PC1** | -0.023612 | 0.016492 | -1.4317 | 0.15233 | -0.055948 | 0.0087254 |
| **PC2** | 0.14124 | 0.027946 | 5.054 | 4.6064e-07 | 0.086442 | 0.19603 |
| **PC3** | 0.039578 | 0.038862 | 1.0184 | 0.30856 | -0.036623 | 0.11578 |
| **PC4** | -0.0096321 | 0.036344 | -0.26502 | 0.79101 | -0.080897 | 0.061633 |

**Table S7.**
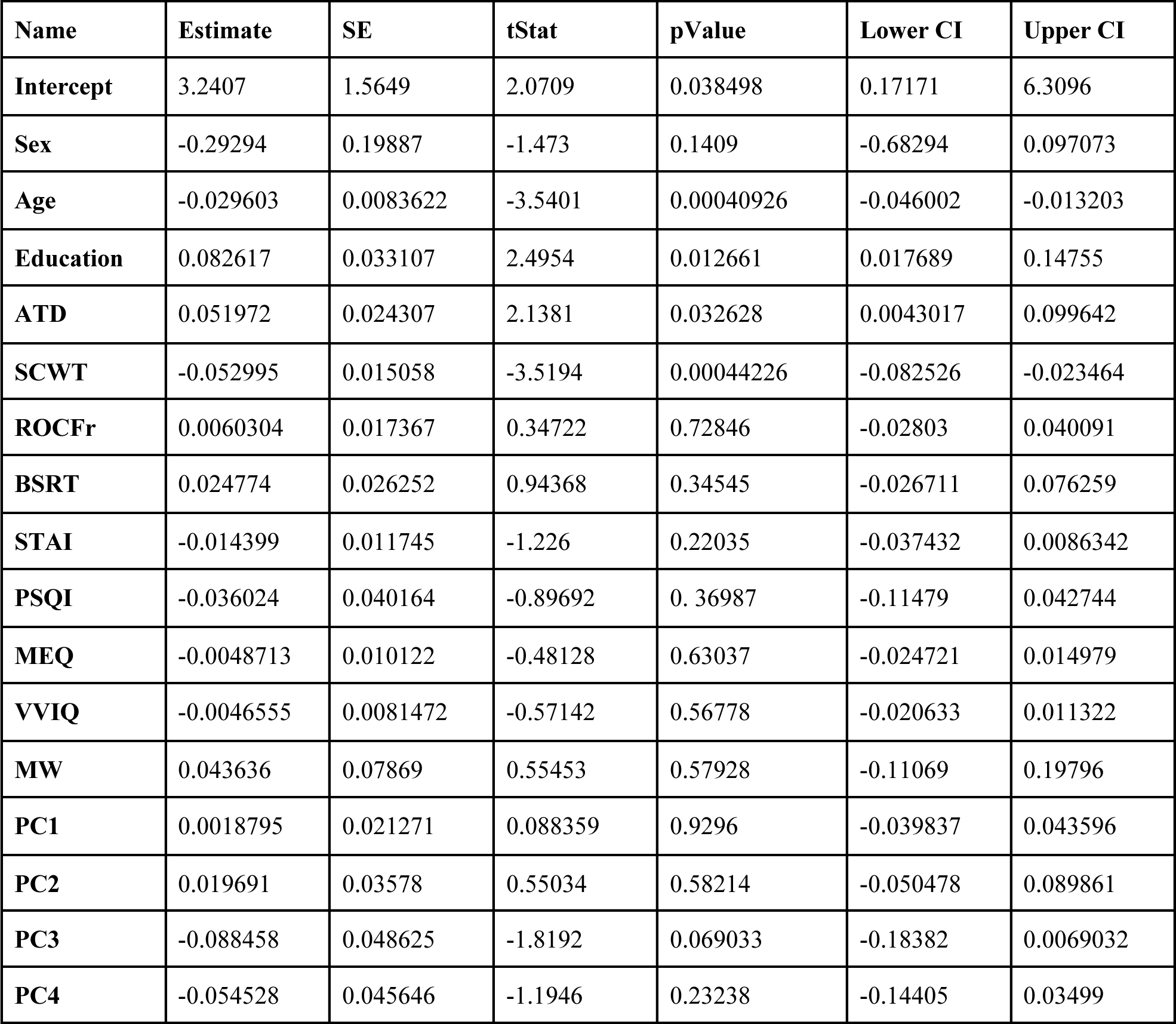
LME results for CD vs. WD (df = 1981). For each predictor, we reported the estimate, standard error (SE), t statistics, p-value, and lower and upper bounds of the effect 95% confidence interval.

| Name | Estimate | SE | tStat | pValue | Lower CI | Upper CI |
| --- | --- | --- | --- | --- | --- | --- |
| **Intercept** | 3.2407 | 1.5649 | 2.0709 | 0.038498 | 0.17171 | 6.3096 |
| **Sex** | -0.29294 | 0.19887 | -1.473 | 0.1409 | -0.68294 | 0.097073 |
| **Age** | -0.029603 | 0.0083622 | -3.5401 | 0.00040926 | -0.046002 | -0.013203 |
| **Education** | 0.082617 | 0.033107 | 2.4954 | 0.012661 | 0.017689 | 0.14755 |
| **ATD** | 0.051972 | 0.024307 | 2.1381 | 0.032628 | 0.0043017 | 0.099642 |
| **SCWT** | -0.052995 | 0.015058 | -3.5194 | 0.00044226 | -0.082526 | -0.023464 |
| **ROCFr** | 0.0060304 | 0.017367 | 0.34722 | 0.72846 | -0.02803 | 0.040091 |
| **BSRT** | 0.024774 | 0.026252 | 0.94368 | 0.34545 | -0.026711 | 0.076259 |
| **STAI** | -0.014399 | 0.011745 | -1.226 | 0.22035 | -0.037432 | 0.0086342 |
| **PSQI** | -0.036024 | 0.040164 | -0.89692 | 0.36987 | -0.11479 | 0.042744 |
| **MEQ** | -0.0048713 | 0.010122 | -0.48128 | 0.63037 | -0.024721 | 0.014979 |
| **VVIQ** | -0.0046555 | 0.0081472 | -0.57142 | 0.56778 | -0.020633 | 0.011322 |
| **MW** | 0.043636 | 0.07869 | 0.55453 | 0.57928 | -0.11069 | 0.19796 |
| **PC1** | 0.0018795 | 0.021271 | 0.088359 | 0.9296 | -0.039837 | 0.043596 |
| **PC2** | 0.019691 | 0.03578 | 0.55034 | 0.58214 | -0.050478 | 0.089861 |
| **PC3** | -0.088458 | 0.048625 | -1.8192 | 0.069033 | -0.18382 | 0.0069032 |
| **PC4** | -0.054528 | 0.045646 | -1.1946 | 0.23238 | -0.14405 | 0.03499 |

## References

[1] L. Salvesen, E. Capriglia, M. Dresler, and G. Bernardi, ‘Influencing dreams through sensory stimulation: A systematic review’, Sleep Med. Rev., vol. 74, p. 101908, 2024, doi: 10.1016/j.smrv.2024.101908.

[2] Y. Nir and G. Tononi, ‘Dreaming and the brain: from phenomenology to neurophysiology’, Trends Cogn. Sci., vol. 14, no. 2, pp. 88–100, 2010, doi: 10.1016/j.tics.2009.12.001.

[3] F. Siclari, K. Valli, and I. Arnulf, ‘Dreams and nightmares in healthy adults and in patients with sleep and neurological disorders’, Lancet Neurol., vol. 19, no. 10, pp. 849–859, 2020, doi: 10.1016/S1474-4422(20)30275-1.

[4] K. Valli, B. Frauscher, V. Gschliesser, E. Wolf, T. Falkenstetter, S. V. Schoenwald, … and B. Hoegl, ‘Can observers link dream content to behaviours in rapid eye movement sleep behaviour disorder? A cross-sectional experimental pilot study: Dream content and behavioural manifestations in RBD’, J. Sleep Res., vol. 21, no. 1, pp. 21–29, 2012, doi: 10.1111/j.1365-2869.2011.00938.x.

[5] I. Arnulf, ‘Dream imagery, rapid eye movement sleep behavior disorder, and hallucinations: Dreaming, visual hallucinations and RBD’, Sleep Biol. Rhythms, vol. 11, pp. 15–20, 2013, doi: 10.1111/j.1479-8425.2012.00546.x.

[6] C. Koch, M. Massimini, M. Boly, and G. Tononi, ‘Neural correlates of consciousness: progress and problems’, Nat. Rev. Neurosci., vol. 17, no. 5, pp. 307–321, 2016, doi: 10.1038/nrn.2016.22.

[7] G. Tononi, M. Boly, and C. Cirelli, ‘Consciousness and sleep’, Neuron, pp. S0896-6273(24)00272–1, 2024, doi: 10.1016/j.neuron.2024.04.011.

[8] E. Aserinsky and N. Kleitman, ‘Regularly occurring periods of eye motility, and concomitant phenomena, during sleep’, Science, vol. 118, pp. 273–274, 1953, doi: 10.1126/science.118.3062.273.

[9] W. Dement and N. Kleitman, ‘Cyclic variations in EEG during sleep and their relation to eye movements, body motility, and dreaming’, Electroencephalogr. Clin. Neurophysiol., vol. 9, no. 4, pp. 673–690, 1957, doi: 10.1016/0013-4694(57)90088-3.

[10] F. Siclari, J. J. LaRocque, B. R. Postle, and G. Tononi, ‘Assessing sleep consciousness within subjects using a serial awakening paradigm’, Front. Psychol., vol. 4, 2013, doi: 10.3389/fpsyg.2013.00542.

[11] G. Nemeth, ‘The route to recall a dream: theoretical considerations and methodological implications’, Psychol. Res., vol. 87, no. 4, pp. 964–987, 2023, doi: 10.1007/s00426-022-01722-7.

[12] E. Fränkl et al., ‘How our Dreams Changed During the COVID-19 Pandemic: Effects and Correlates of Dream Recall Frequency - a Multinational Study on 19,355 Adults’, Nat. Sci. Sleep, vol. 13, pp. 1573–1591, 2021, doi: 10.2147/NSS.S324142.

[13] P. Fazekas, G. Nemeth, and M. Overgaard, ‘White dreams are made of colours: What studying contentless dreams can teach about the neural basis of dreaming and conscious experiences’, Sleep Med. Rev., vol. 43, pp. 84–91, 2019, doi: 10.1016/j.smrv.2018.10.005.

[14] J. Windt, ‘Consciousness in sleep: How findings from sleep and dream research challenge our understanding of sleep, waking, and consciousness’, Philos. Compass, vol. 15, 2020, doi: 10.1111/phc3.12661.

[15] M. Blagrove and E. F. Pace-Schott, ‘Trait And Neurobiological Correlates Of Individual Differences In Dream Recall And Dream Content’, in International Review of Neurobiology, vol. 92, A. Clow and P. McNamara, Eds., Academic Press, 2010, pp. 155–180. doi: 10.1016/S0074-7742(10)92008-4.

[16] S. F. Butler and R. Watson, ‘Individual differences in memory for dreams: The role of cognitive skills’, Percept. Mot. Skills, vol. 61, no. 3, Pt 1, pp. 823–828, 1985, doi: 10.2466/pms.1985.61.3.823.

[17] J.-B. Eichenlaub, A. Nicolas, J. Daltrozzo, J. Redouté, N. Costes, and P. Ruby, ‘Resting Brain Activity Varies with Dream Recall Frequency Between Subjects’, Neuropsychopharmacology, vol. 39, no. 7, Art. no. 7, 2014, doi: 10.1038/npp.2014.6.

[18] D. B. Cohen, ‘Dream Recall and Short-Term Memory’, Percept. Mot. Skills, vol. 33, no. 3, pp. 867–871, 1971, doi: 10.2466/pms.1971.33.3.867.

[19] M. Schredl, ‘Dream recall frequency in a representative German sample’, Percept. Mot. Skills, vol. 106, no. 3, pp. 699–702, 2008, doi: 10.2466/pms.106.3.699-702.

[20] M. Schredl, ‘Explaining the Gender Difference in Dream Recall Frequency’, Dreaming, vol. 20, pp. 96–106, 2010, doi: 10.1037/a0019392.

[21] P. M. Ruby, ‘The Neural Correlates of Dreaming Have Not Been Identified Yet. Commentary on “The Neural Correlates of Dreaming. Nat Neurosci. 2017”’, Front. Neurosci., vol. 14, 2020.

[22] M. Schredl, Researching Dreams: The Fundamentals. Cham: Springer International Publishing, 2018. doi: 10.1007/978-3-319-95453-0.

[23] K. Christoff, Z. C. Irving, K. C. R. Fox, R. N. Spreng, and J. R. Andrews-Hanna, ‘Mind-wandering as spontaneous thought: a dynamic framework’, Nat. Rev. Neurosci., vol. 17, no. 11, pp. 718–731, 2016, doi: 10.1038/nrn.2016.113.

[24] K. C. R. Fox, S. Nijeboer, E. Solomonova, G. W. Domhoff, and K. Christoff, ‘Dreaming as mind wandering: evidence from functional neuroimaging and first-person content reports’, Front. Hum. Neurosci., vol. 7, p. 412, 2013, doi: 10.3389/fnhum.2013.00412.

[25] G. W. Domhoff, The scientific study of dreams: Neural networks, cognitive development, and content analysis. in The scientific study of dreams: Neural networks, cognitive development, and content analysis. Washington, DC, US: American Psychological Association, 2003, pp. ix, 209. doi: 10.1037/10463-000.

[26] J. Smallwood and J. W. Schooler, ‘The science of mind wandering: empirically navigating the stream of consciousness’, Annu. Rev. Psychol., vol. 66, pp. 487–518, 2015, doi: 10.1146/annurev-psych-010814-015331.

[27] P. Ruby, C. Blochet, J. Eichenlaub, O. Bertrand, D. Morlet, and A. Bidet-Caulet, ‘Alpha reactivity to first names differs in subjects with high and low dream recall frequency’, Front. Psychol., vol. 4, p. 419, 2013, doi: 10.3389/fpsyg.2013.00419.

[28] M. E. Raichle and A. Z. Snyder, ‘A default mode of brain function: a brief history of an evolving idea’, NeuroImage, vol. 37, no. 4, pp. 1083–1090; discussion 1097-1099, 2007, doi: 10.1016/j.neuroimage.2007.02.041.

[29] R. L. Buckner, J. R. Andrews-Hanna, and D. L. Schacter, ‘The brain’s default network: anatomy, function, and relevance to disease’, Ann. N. Y. Acad. Sci., vol. 1124, pp. 1–38, 2008, doi: 10.1196/annals.1440.011.

[30] M. F. Mason, M. I. Norton, J. D. Van Horn, D. M. Wegner, S. T. Grafton, and C. N. Macrae, ‘Wandering Minds: The Default Network and Stimulus-Independent Thought’, Science, vol. 315, no. 5810, pp. 393–395, 2007, doi: 10.1126/science.1131295.

[31] F. Siclari, G. Bernardi, J. Cataldi, and G. Tononi, ‘Dreaming in NREM Sleep: A High-Density EEG Study of Slow Waves and Spindles’, J. Neurosci., vol. 38, no. 43, pp. 9175–9185, 2018, doi: 10.1523/JNEUROSCI.0855-18.2018.

[32] F. Siclari et al., ‘The neural correlates of dreaming’, Nat. Neurosci., vol. 20, no. 6, pp. 872–878, 2017, doi: 10.1038/nn.4545.

[33] J. Zhang and E. J. Wamsley, ‘EEG predictors of dreaming outside of REM sleep’, Psychophysiology, vol. 56, no. 7, p. e13368, 2019, doi: 10.1111/psyp.13368.

[34] S. Scarpelli et al., ‘Predicting Dream Recall: EEG Activation During NREM Sleep or Shared Mechanisms with Wakefulness?’, Brain Topogr., vol. 30, no. 5, pp. 629–638, 2017, doi: 10.1007/s10548-017-0563-1.

[35] M. Massimini, F. Ferrarelli, R. Huber, S. K. Esser, H. Singh, and G. Tononi, ‘Breakdown of Cortical Effective Connectivity During Sleep’, Science, vol. 309, no. 5744, pp. 2228–2232, 2005, doi: 10.1126/science.1117256.

[36] A. Pigorini et al., ‘Bistability breaks-off deterministic responses to intracortical stimulation during non-REM sleep’, NeuroImage, vol. 112, pp. 105–113, 2015, doi: 10.1016/j.neuroimage.2015.02.056.

[37] B. A. Riedner et al., ‘Sleep Homeostasis and Cortical Synchronization: III. A High-Density EEG Study of Sleep Slow Waves in Humans’, Sleep, vol. 30, no. 12, pp. 1643–1657, 2007, doi: 10.1093/sleep/30.12.1643.

[38] D. B. Cohen and G. Wolfe, ‘Dream recall and repression: evidence for an anternative hypothesis’, J. Consult. Clin. Psychol., vol. 41, no. 3, pp. 349–355, 1973, doi: 10.1037/h0035333.

[39] T. Nielsen, ‘Variations in Dream Recall Frequency and Dream Theme Diversity by Age and Sex’, Front. Neurol., vol. 3, p. 106, 2012, doi: 10.3389/fneur.2012.00106.

[40] M. Schredl and I. Reinhard, ‘Gender differences in dream recall: a meta-analysis’, J. Sleep Res., vol. 17, no. 2, pp. 125–131, 2008, doi: 10.1111/j.1365-2869.2008.00626.x.

[41] K. Bulkeley and M. Schredl, ‘Attitudes towards dreaming: Effects of socio-demographic and religious variables in an American sample’, vol. 12, no. 1, p. 7, 2019.

[42] S. Blain, A. de la Chapelle, A. Caclin, A. Bidet-Caulet, and P. Ruby, ‘Dream recall frequency is associated with attention rather than with working memory abilities’, J. Sleep Res., vol. 31, no. 5, p. e13557, 2022, doi: 10.1111/jsr.13557.

[43] C. Sikaras et al., ‘The Mediating Role of Depression and of State Anxiety οn the Relationship between Trait Anxiety and Fatigue in Nurses during the Pandemic Crisis’, Healthcare, vol. 11, no. 3, Art. no. 3, 2023, doi: 10.3390/healthcare11030367.

[44] D. F. Marks, ‘Visual imagery differences in the recall of pictures’, Br. J. Psychol. Lond. Engl. 1953, vol. 64, no. 1, pp. 17–24, 1973, doi: 10.1111/j.2044-8295.1973.tb01322.x.

[45] J. S. A. Carriere, P. Seli, and D. Smilek, ‘Wandering in both mind and body: individual differences in mind wandering and inattention predict fidgeting’, Can. J. Exp. Psychol. Rev. Can. Psychol. Exp., vol. 67, no. 1, pp. 19–31, 2013, doi: 10.1037/a0031438.

[46] J. Backhaus, K. Junghanns, A. Broocks, D. Riemann, and F. Hohagen, ‘Test-retest reliability and validity of the Pittsburgh Sleep Quality Index in primary insomnia’, J. Psychosom. Res., vol. 53, no. 3, pp. 737–740, 2002, doi: 10.1016/s0022-3999(02)00330-6.

[47] J. A. Horne and O. Ostberg, ‘A self-assessment questionnaire to determine morningness-eveningness in human circadian rhythms’, Int. J. Chronobiol., vol. 4, no. 2, pp. 97–110, 1976.

[48] M. Schredl, ‘Questionnaires and diaries as research instruments in dream research: Methodological issues’, Dreaming, vol. 12, no. 1, pp. 17–26, 2002, doi: 10.1023/A:1013890421674.

[49] F. Scarpina and S. Tagini, ‘The Stroop Color and Word Test’, Front. Psychol., vol. 8, p. 557, 2017, doi: 10.3389/fpsyg.2017.00557.

[50] M. D. Horner, G. Teichner, K. B. Kortte, and R. T. Harvey, ‘Construct validity of the Babcock Story Recall Test’, Appl. Neuropsychol., vol. 9, no. 2, pp. 114–116, 2002, doi: 10.1207/S15324826AN0902_7.

[51] M.-S. Shin, S.-Y. Park, S.-R. Park, S.-H. Seol, and J. S. Kwon, ‘Clinical and empirical applications of the Rey-Osterrieth Complex Figure Test’, Nat. Protoc., vol. 1, no. 2, pp. 892–899, 2006, doi: 10.1038/nprot.2006.115.

[52] D. Fekedulegn, M. E. Andrew, M. Shi, J. M. Violanti, S. Knox, and K. E. Innes, ‘Actigraphy-Based Assessment of Sleep Parameters’, Ann. Work Expo. Health, vol. 64, no. 4, pp. 350–367, 2020, doi: 10.1093/annweh/wxaa007.

[53] P. J. Arnal et al., ‘The Dreem Headband compared to polysomnography for electroencephalographic signal acquisition and sleep staging’, Sleep, vol. 43, no. 11, p. zsaa097, 2020, doi: 10.1093/sleep/zsaa097.

[54] Y. Benjamini and Y. Hochberg, ‘Controlling the False Discovery Rate: A Practical and Powerful Approach to Multiple Testing’, J. R. Stat. Soc. Ser. B Methodol., vol. 57, no. 1, pp. 289–300, 1995, doi: 10.1111/j.2517-6161.1995.tb02031.x.

